# Using Genome-Wide Association Study to Identify Genes and Pathways associated with Hypersensitivity Pneumonitis

**DOI:** 10.1101/2020.05.27.118539

**Authors:** Ling Wang, Melissa May Millerick, Kenneth D. Rosenman, Yuehua Cui, Bruce Uhal, Jianrong Wang, John Gerlach

## Abstract

**Background:** Hypersensitivity Pneumonitis (HP) is an interstitial lung disease caused by an immune response to the inhalation of antigens. Since only a small proportion of individuals exposed to HP-related antigens develop the disease, a genetic variation may play a role in disease development.

**Methods:** In this small-scale study, 24 patients diagnosed with HP were matched with control group who shared the patient’s environment and were exposed to the same HP-associated antigens. Logistic regression was employed to identify Single-Nucleotide Polymorphisms (SNPs) associated with HP. Next genes associated with HP were identified using sequence kernel association test (SKAT) analysis. Last, Kyoto Encyclopedia of Genes and Genomes (KEGG) and Gene Oncology (GO) enrichment analysis were employed to find HP signaling pathways using SNPs coded on genes and on non-coding genes, respectively.

**Results:** Given the small sample size, no single SNPs or genes were identified to be significantly associated with HP after adjustment for multiple testing. After P-value adjustment, the KEGG and GO pathway enrichment analysis identified 11 and 20 significant pathways respectively using SNPs coded on genes. Among these pathways, Cell cycle, Proteasome and Base excision repair had previously reported to be associated with lung function.

**Conclusion:** This is the first GWAS study identifying genetic factors associated with HP. Although no significant associations at SNPs/gene level were identified, there were significant pathways that are identified associated with HP which need further investigation in large cohorts.

## Introduction

Hypersensitivity pneumonitis (HP) is an interstitial lung disease resulting from complex interactions between antigen exposures and alleles of many genes^1–4^. HP development depends on the type, intensity, and duration of exposure to the inciting agent, host susceptibility, the site of interaction within the respiratory system, and the resulting level of dysregulation of the cellular and humoral immune response over time^5^. In the United States there is a significant increasing trend in overall age-adjusted mortality for HP^6^, with HP mortality in farm workers 10-50% more than expected^7^.

Only a small proportion of exposed individuals develop HP indicating there are likely to be genetic factors involved in the development of the disease. It has been found by previous researchers that genes of the major histocompatibility complex (MHC) and the tumor necrosis factor alpha (TNF-α) were associated with HP^4,5,8^. The expression of MHC class II molecules is required for activation of T lymphocytes cells during immune response triggering. TNF-α is a pro-inflammatory cytokine produced primarily in the Th1-like microenvironment as occurs in HP. Recent review by Kiszałkiewicz et al. suggested several signaling pathways (e.g., TNF-α/NFκβ, TGF-β/SMAD, Wnt-β-catenin) are activated in idiopathic pulmonary fibrosis (IPF) and HP^9^. Ley et al. (2019) found that rare variants in telomere-related genes are significantly associated with HP and reduced transplant-free survival in HP patients^10^. However, all previous HP genetic studies have concentrated on pre-selected genes, and thus have a limited ability to comprehensively identify genetic factors associated with HP. Furthermore, previous studies simply chose healthy individuals as controls, rather than controls matched with similar environmental exposure.

In this study, we used a matched-pair design with each HP patient (case) matched with a control subject (a family member or co-worker who shared the same environment as the patient) and employed genome-wide association (GWA) methods to identify genes associated with HP. To overcome the problem of low power due to small sample size in this pilot study, we grouped functionally HP-related genes into biological modules using pathway analysis. We hypothesize that a complex disease such as HP is likely to have causal genes exerting their effects through small perturbations in biological pathways. Rather than turning proteins on or off, they may subtly alter the amount of proteins produced. Furthermore, because the majority of the human genome are not protein-coding sequences^11^, large numbers of significant SNPs from GWAS studies are located in non-coding regions. The non-coding GWAS SNPs have been shown to be enriched in regulatory elements which can control tissue-specific gene expression^12^. Therefore, we also investigated non-coding SNPs located in regulatory elements to improve the identification of associated pathways in HP.

## Materials and Methods

### Study Population

Twenty-four HP patients who met standard diagnostic criteria for HP^13^ were recruited from the pulmonary practices associated with three academic/tertiary hospitals in Michigan. This study received IRB approval, and informed consent was obtained from all subjects.

Each HP case was asked to identify one ‘control’. If the suspected exposure of interest was in the home, the case was asked to select an adult family member as his/her corresponding control. If the suspected exposure of interest was from the workplace, then the case was asked to identify a co-worker as his/her control.

### Genomic DNA isolation

At least 2 mL of whole blood was collected in an EDTA blood tube (Beckton Dickinson) from both HP patients and controls for genetic analysis. DNA was extracted from 200uL of the whole blood with concentration > 50ng/ul (QiaSymphony DNA extraction kit, Qiagen), and then quantified (PicoGreen fluorometric dye, ThermoFisher). Samples were split and stored in -20°C to -80°C freezers. The laboratory performing the genotyping was blinded as to the patient/control status.

### Genotyping

Two hundred microliters of DNA from each HP patient and control were applied to the Illumina Omni 2.5 GWAS array (https://www.illumina.com/products/by-type/microarray-kits/infinium-omni25-8.html), which contains 2.3 million common and rare SNPs curated from the 1000 Genomes Project for diverse world populations. A total of 2,382,209 SNPs were genotyped for 48 samples in the analysis. Genotype frequencies were tested by Hardy-Weinberg equilibrium.

### Quality Control

Quality Control (QC) was done in two steps: 1) the missingness threshold for SNPs was set to 0.2 and 28,610 SNPs were excluded; 2) two thresholds for the Hardy-Weinberg equilibrium were used: 10^−10^ was the P-value threshold for excluding SNPs in HP cases, and 10^−6^ was the P-value threshold for excluding SNPs in controls^14^. Ten SNPs in HP cases and 10 SNPs in controls failed the test. After QC, there were 2,353,579 SNPs left. We didn’t exclude rare variants in this study because rare variants were found to be associated with HP significantly^10^.

### Mapping SNP IDs to Genes

The SNP IDs were mapped to gene IDs using the “biomaRT” R package, which provides an interface to a growing collection of databases implementing the BioMart software suite^15^. The data set “hsapiens_snp” in “snp” biomart were used to make queries of SNPs IDs that matched to gene IDs.

### Statistical Methods

We first used the standard approach in genome-wide association studies (GWAS), which was to test the association between HP status and SNPs using logistic regression controlling for two principle components. Based on the Bonferroni correction, we used a significance threshold of 5×10^−8^ as commonly used in traditional GWAS.

Next, instead of testing each SNP individually, we employed the sequence kernel association test (SKAT)^12,13^ to identify significant genes associated with HP. SKAT is a flexible, computationally efficient, regression approach that tests for association between variants in a region (both common and rare) and a dichotomous or continuous phenotype while adjusting for covariates, such as principal components to account for population stratification. Specifically, assume *y*_*i*_ indicating HP status for individual *i* and consider the following generalized linear model: 

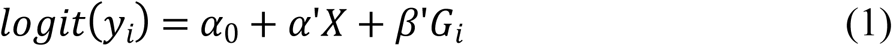

where *X* are the covariates and *G*_*i*_ contains *G*_*ij*_, the genotype variant *j* for individual *i*. SKAT assumes that regression coefficients *β*_*j*_ for variant *G*_*ij*_ follow a distribution of mean 0, and variance 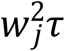 and test the hypothesis *τ* = 0 using a variance-component score test. The test statistic is defined as 

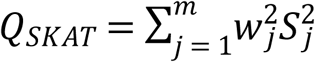

where 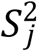 is the score statistics of genetic variants *G*_*ij*_ in model (1). *w*_*j*_ is a weight assigned to 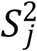. It is a random number drawn from *beta*(*MAF*_*i*_, 1,25) distribution in our study, where *beta* is a beta density function and *MAF*_*i*_ is a minor allele frequency (MAF) of SNP *i*. This weight *w*_*j*_ allows for increasing the weight of rare variants while still putting decent nonzero weights for common variants^16^. *Q*_*SKAT*_ asymptotically follows a mixture chi-square distribution. SKAT analyses were performed using the SKAT package in R (https://cran.r-project.org/web/packages/SKAT/index.html).

To further reduce the complexity of analysis while simultaneously providing greater explanatory power, we used both Kyoto Encyclopedia of Genes and Genomes (KEGG)^17^ and Gene Ontology (GO)^18^ enrichment pathway to find groups of genes that function in the same pathways. For significant genes identified by SKAT analysis (P<0.05), an active subnetwork is defined as a group of inter-connected genes in a protein-protein interaction network (PIN) that mostly consists of significant genes^19^. The relations among genes were added to the PIN as undirected links, removing any duplicate interactions. Pathway enrichment analyses were preformed using the “pathfinder” package in R (https://cran.r-project.org/web/packages/pathfindR/index.html).

Since more than 95% of the human genome is non-coding, most significant SNPs are located in non-coding regions. In traditional pathway analysis, non-coding SNPs have limited power to identify disease-associated genes, due to the missing information of specific target genes linked to non-coding SNPs^20^. We overcame this problem by linking non-coding SNPs to their potential target genes, whose expression may be dysregulated by these non-coding SNPs. In this way, we substantially expanded the set of candidate HP-associated genes. We used the genome-wide regulatory element annotation from Roadmap Epigenomics project and ENCODE project^12,21^ and identified the subset of significant SNPs that are located in regulatory elements. We further calculated the activity correlations between specific regulatory elements and genes across multiple cell-types and tissues from ENCODE and Roadmap Epigenomics dataset. For regulatory element activity, we used the epigenetic features that have been shown to represent the functional activity levels of regulatory elements^22^, i.e. H3K4me1 and H3K27me3. For gene activity, we used the RNA-seq data to represent gene expression levels. We also randomly shuffled the matrix of gene expression 1,000 times and calculated the correlations for each pair of regulatory elements and genes. Based on the null distribution from the 1,000 shuffled sets, we calculated empirical p-values for each pair of regulatory element and gene. The genes with significant activity correlations (False Discovery Rate (FDR) < 0.05, Benjamini-Hochberg correction) were considered as target genes for specific regulatory elements. To further reduce the potential false positives, we only considered significantly correlated genes whose promoters are located within +/-1 Mb from the regulatory element as the final candidate target gene. we further carried out pathway enrichment analysis^23,24^ on the gene sets and identified biological pathways that may be disrupted by non-coding HP-SNPs.

## Results

The demographic information for HP patients and controls are shown in Table 1. There are no significant differences in basic demographics between HP patients and controls.

**Table 1.**
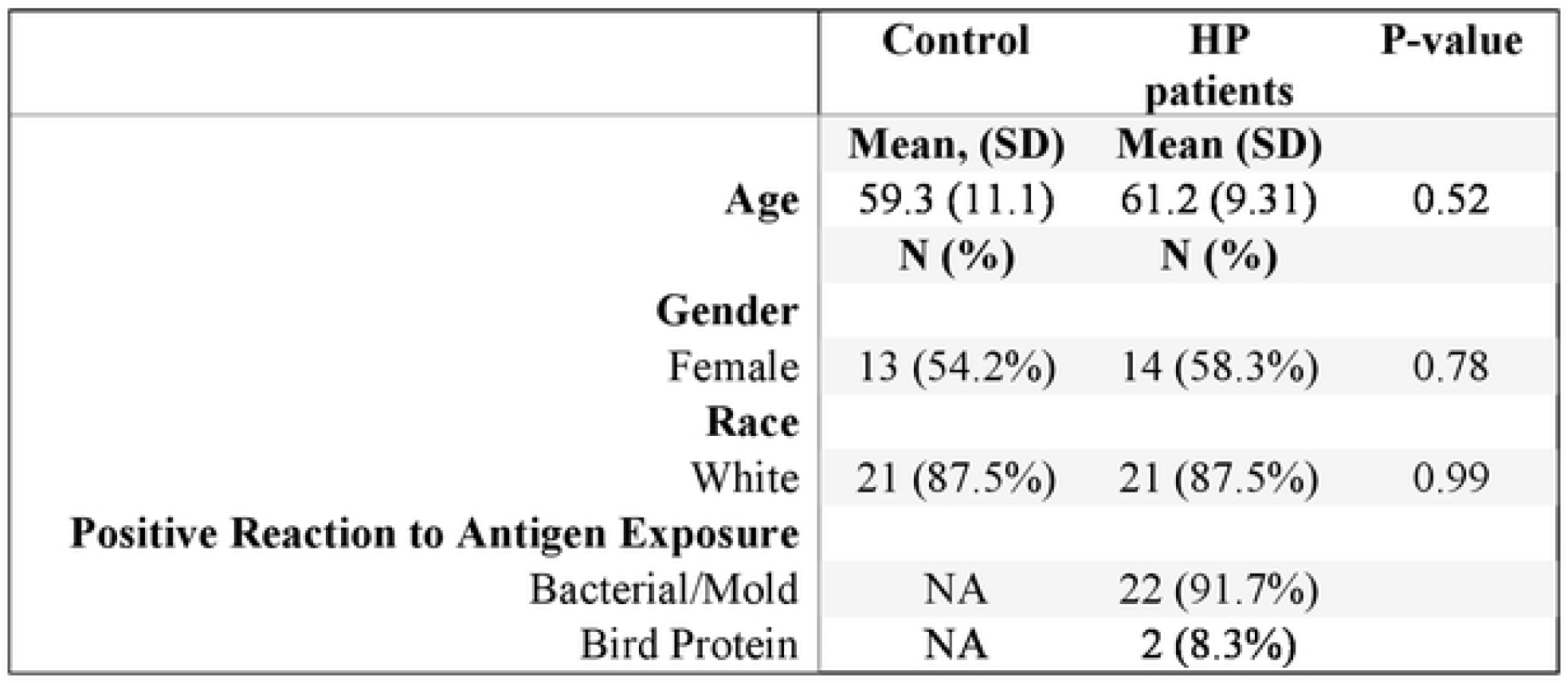
Demographics of HP patients and matched controls

### Single-SNP Tests

Figure 1 shows the Manhattan Plot for association between SNPs with HP disease using logistic regression controlling for the first two principle components. After Bonferroni correct, none of the SNPs reached statistical significance (P<5×10^−8^).

**Figure 1.**
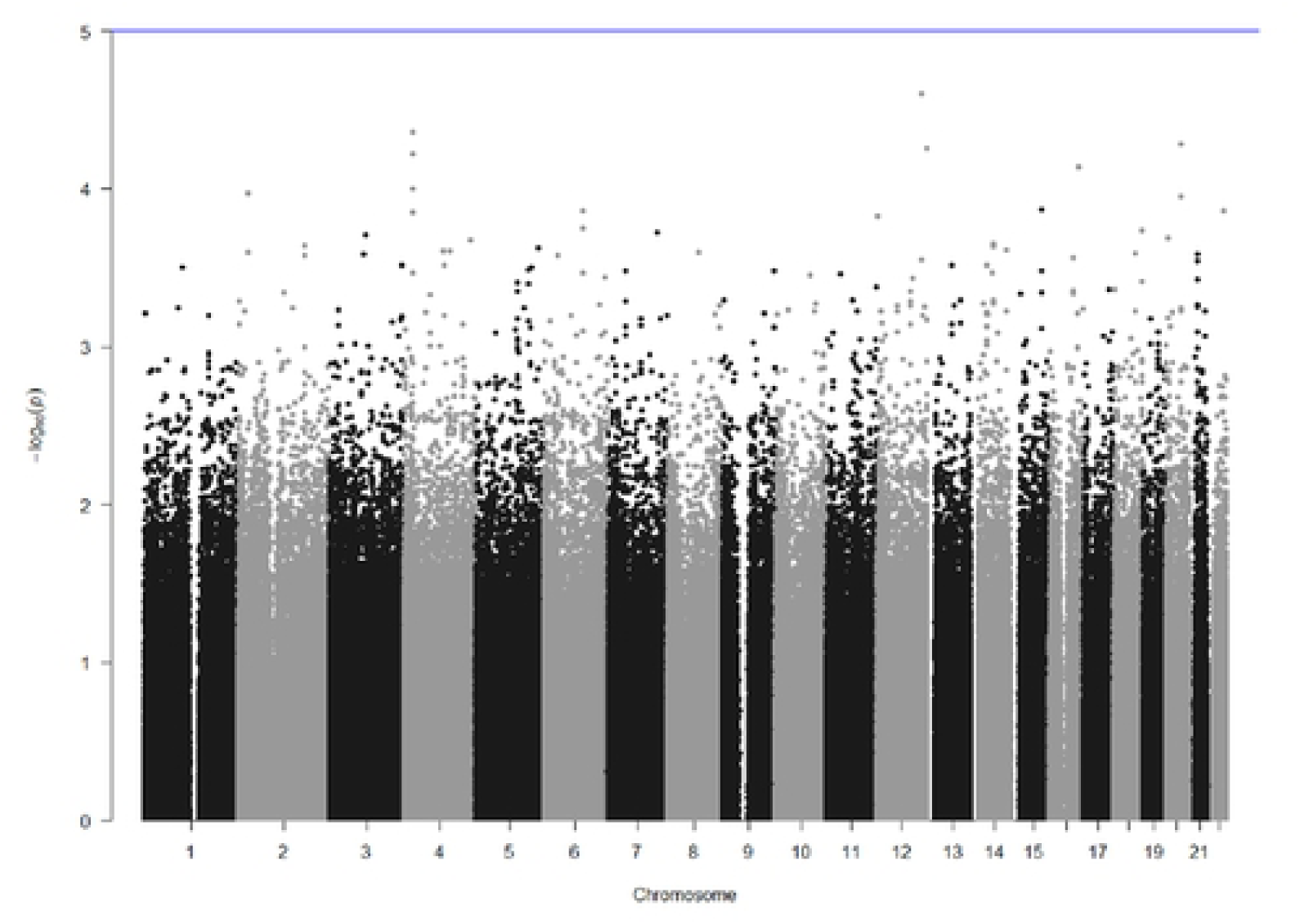
Manhattan Plot of logistic regression results by single SNP analysis, where the X-axis shows the chromosomal position and the y-axis shows the −log (P-value) for each SNP in the genome-wide analysis. The blue line shows suggested genome-wide significance (P<10^−5^).

### Gene Tests

There were 944,748 SNPs on 36,563 genes identified in our sample. The association between genes and HP disease were analyzed by SKAT controlling for the first 2 PCs. Table 2 shows the top 25 genes with the smallest P-values. After Benjamini-Hochberg correction^25^, there were no genes significantly associated with HP at α=0.05.

**Table 2.** Top 25 Genes with the smallest P-values associated with HP identified by SKAT analysis

| Gene IDs | Chromosome | Number of SNPs | Start BP | P-value |
| --- | --- | --- | --- | --- |
| ***FEZ2*** | 2 | 89 | 36621826 | 8.79E-06 |
| ***CELF1*** | 11 | 23 | 47514250 | 2.18E-05 |
| ***OR4C9P*** | 11 | 1 | 48464325 | 7.96E-05 |
| ***H6PD*** | 1 | 35 | 9265617 | 0.000151 |
| ***EMB*** | 5 | 13 | 50439676 | 0.000199 |
| ***IL5*** | 5 | 4 | 1.33E+08 | 0.000215 |
| ***VRTN*** | 14 | 24 | 74350715 | 0.00022 |
| ***RPA2P1*** | 14 | 3 | 46996797 | 0.000264 |
| ***LINC01788*** | 1 | 31 | 70744035 | 0.000267 |
| ***PSMC1P10*** | 2 | 1 | 17385565 | 0.000347 |
| ***NDUFS3*** | 11 | 3 | 47573499 | 0.000363 |
| ***NR1H3*** | 11 | 6 | 47268126 | 0.000416 |
| ***SPINK1*** | 5 | 6 | 1.48E+08 | 0.000432 |
| ***DDB2*** | 11 | 11 | 47216129 | 0.000524 |
| ***ZNF165*** | 6 | 12 | 28083000 | 0.00053 |
| ***OR4X2*** | 11 | 2 | 48245427 | 0.000591 |
| ***RIMS3*** | 1 | 28 | 40632806 | 0.000596 |
| ***OR4X1*** | 11 | 6 | 48264354 | 0.0006 |
| ***ADSSL1*** | 14 | 10 | 1.05E+08 | 0.000693 |
| ***TMEM244*** | 6 | 28 | 1.3E+08 | 0.000799 |
| ***INTU*** | 4 | 23 | 1.28E+08 | 0.000827 |
| ***RASSF4*** | 10 | 29 | 44977730 | 0.000871 |
| ***ELF1*** | 13 | 47 | 40967878 | 0.000901 |
| ***SLC35F2*** | 11 | 79 | 1.08E+08 | 0.001017 |
| ***OPA1*** | 3 | 42 | 1.94E+08 | 0.00109 |

### Pathway analysis

#### Genetic variants coded on genes

To increase the power of the study, we examined groups of genes association with HP using KEGG and GO enrichment analysis. Genes with P-values ≤ 0.05 calculated from SKAT analysis (N=1,520) were used in the pathway enrichment analysis.

Table 3 shows the eleven pathways significantly correlated with HP using KEGG enrichment analysis (FDR <0.05 after Benjamini-Hochberg correction^25^). The top five pathways identified were: 1) Cell cycle (Adjusted P=0.0002) 2) Cellular senescence (Adjusted P=0.0009) 3) Proteasome (Adjusted P=0.0009) 4) Base excision repair (Adjusted P=0.0009) and 5) Ribosome (Adjusted P=0.0035).

**Table 3.** Significant KEGG/ GO pathways identified using SNPs on coded genes

| **KEGG Pathways** | **P-value** | **Adjusted P-value** |
| --- | --- | --- |
| Cell cycle | 4.94E-06 | 0.000193 |
| Protein processing in endoplasmic reticulum | 5.59E-05 | 0.000899 |
| Cellular senescence | 8.13E-05 | 0.000899 |
| p53 signaling pathway | 9.22E-05 | 0.000899 |
| Proteasome | 0.000452 | 0.003522 |
| Viral carcinogenesis | 0.002376 | 0.015447 |
| Notch signaling pathway | 0.004006 | 0.022322 |
| Epstein-Barr virus infection | 0.006571 | 0.029072 |
| Adherens junction | 0.006709 | 0.029072 |
| Hippo signaling pathway - multiple species | 0.008534 | 0.033043 |
| Ubiquitin mediated proteolysis | 0.010338 | 0.033043 |
| Ribosome | 0.011014 | 0.033043 |
| **GO Pathways** |  |  |
| Histone deacetylase binding | 3.32E-10 | 3.15E-08 |
| G2/M transition of mitotic cell cycle | 4.00E-07 | 3.76E-05 |
| Protein deubiquitination | 1.90E-06 | 0.000177 |
| Ciliary basal body-plasma membrane docking | 6.23E-06 | 0.000573 |
| Centriole | 9.36E-06 | 0.000852 |
| Protein-DNA complex | 3.04E-05 | 0.002732 |
| Embryonic digit morphogenesis | 6.67E-05 | 0.005937 |
| Thiol-dependent ubiquitinyl hydrolase activity | 8.29E-05 | 0.007299 |
| Protein polyubiquitination | 0.000109 | 0.009465 |
| Regulation of G2/M transition of mitotic cell cycle | 0.00011 | 0.009465 |
| NF-kappaB binding | 0.000149 | 0.012703 |
| Thiol-dependent ubiquitin-specific protease activity | 0.000182 | 0.015307 |
| Interleukin-1-mediated signaling pathway | 0.000262 | 0.021742 |
| Nucleotide-excision repair, DNA damage recognition | 0.000285 | 0.02336 |
| Nucleotide-excision repair, DNA duplex unwinding | 0.000285 | 0.02336 |
| Transcription-coupled nucleotide-excision repair | 0.000412 | 0.03293 |
| Anaphase-promoting complex-dependent catabolic process | 0.00042 | 0.033143 |
| Regulation of mRNA stability | 0.000441 | 0.034367 |
| Regulation of signal transduction by p53 class mediator | 0.000513 | 0.039463 |
| Global genome nucleotide-excision repair | 0.000579 | 0.043987 |

Table 3 also shows the twenty significant pathways identified by GO enrichment analysis (FDR <0.05 after Benjamini-Hochberg correction^25^). The top five pathways were: 1) Histone deacetylase binding (Adjusted P<0.0001) 2) G2/M transition of mitotic cell cycle (Adjusted P<0.0001) 3) Protein deubiquitination (Adjusted P=0.0002) 4) ciliary basal body-plasma membrane docking (Adjusted P=0.0006) and 5) centriole (Adjusted P=0.0009)

#### Genetic variants not coded on genes

Table 4 shows the significant GO pathways identified using genes regulated by SNPs not coded on genes. However, after adjusting for multiple testing, no significant Go pathways were identified.

**Table 4.** GO Pathways identified using genes regulated by SNPs on non-coding regions

| **GO Pathways** | **P-Value** | **Adjusted P-value** |
| --- | --- | --- |
| Cytoskeleton organization | 0.001 | 0.310 |
| Positive regulation of DNA replication | 0.005 | 0.720 |
| Positive regulation of cell proliferation | 0.007 | 0.660 |
| Positive regulation of insulin receptor signaling pathway | 0.033 | 0.980 |
| Positive regulation of glycolytic process | 0.035 | 0.970 |
| Protein complex assembly | 0.036 | 0.950 |
| Cellular protein metabolic process | 0.037 | 0.930 |
| Positive regulation of transcription from RNA polymerase II promoter | 0.038 | 0.910 |
| Response to muscle activity | 0.038 | 0.880 |
| Positive regulation of glycogen biosynthetic process | 0.038 | 0.880 |
| Intermediate filament cytoskeleton organization | 0.038 | 0.880 |
| Positive regulation of NF-kappaB transcription factor activity | 0.046 | 0.900 |
| Negative regulation of phosphorylation | 0.050 | 0.900 |

## Discussion

This study is the first GWAS study to identify genetic factors associated with HP using matched antigen exposure pairs. The genetic variants were examined at the SNP, gene and pathway levels for their association with HP. Traditional GWAS methods suffer from low power when sample size is not large enough. HP is a rare disease and it is hard to recruit large number of HP patients to gain enough power using traditional GWAS methods. By grouping SNPs on the genes and testing each gene as a unit, we are reducing the number of hypotheses being tested and thus relaxing the stringent conditions for reaching genome-wide significance. Furthermore, if there are multiple independent causal SNPs on the same gene, by considering their joint effects, we will have more power to detect their joint activity. The advantage of SKAT analysis is that it overcomes the traditional GWAS’s problem of having low power for rare variants^26^. Rare genetic variants may play key roles in HP development^10^.

Some genes with small P-values identified in our study were also found in previous studies to be associated with HP or related lung diseases. For example, we identified the ***IL5*** gene as the top gene with the smallest P-values associated with HP. The ***IL5*** gene encodes a cytokine that acts as a growth and differentiation factor for both B cells and eosinophils. This cytokine functions by binding to its receptor, which is a heterodimer, whose beta subunit is shared with the receptors for interleukin 3 (***IL3***) and colony stimulating factor 2 (CSF2/GM-CSF). Walker et al. (1994) analyzed cytokine pattern present in BAL fluid of sarcoidosis patients and demonstrated an increase level of IL5^27^. In addition, ***CELF1*** gene was associated with GU-rich elements (GREs) containing mRNAs, that encoded proteins involves in apoptosis, cell proliferation and cell motility^28^. The ***Lef1*** gene was also identified by Konigshoff et al. (2008) as significantly expressed in Idiopathic Pulmonary Fibrosis (IPF)^29^. Our findings related to ***Lef1*** thus suggest that some signaling pathways known to be activated in IPF may also be involved in HP pathogenesis, but the exact roles of these paths remain to be elucidated. In addition, ***EMB*** gene encodes a transmembrane glycoprotein that is a member of the immunoglobulin superfamily. The encoded protein may be involved in cell growth and development by mediating interactions between the cell and extracellular matrix.

The significant signaling pathways identified in this study were also found in previous studies. For example, the cell cycle pathway was found to be significantly associated with HP (Adjusted P=0.0002). The histopathology of human lung fibrosis has classically described “hyperplastic epithelium”, which is now known to be comprised of both rapidly proliferating and apoptotic epithelial cells^30^. In addition, the histopathology of IPF includes the presence of “fibroblastic foci” comprised of fibroblasts that are proliferating, differentiating into myofibroblasts and depositing collagens^31^.

A proteasome pathway was also associated with HP disease (Adjusted P=0.009) in our study. Proteasomal degradation of misfolded proteins is a known consequence of mutations that result in misfolding of the encoded protein. The proteome of the cell relies on selective proteolysis of cellular proteins. The 26S proteasome is a multi-subunit proteolytic unit of the cell that can lead to preferential cleavage of proteins into peptides that can be presented through MHC/HLA molecules on the surface of the cell to T cell receptors. Continued investigation of the gene set indicated by the KEGG and GO analysis with a larger pool of affected and control samples is warranted as this may provide insight into a unique set of presented peptides that could then be targeted for therapy.

Another unique contribution of our study was that we linked non-coding SNPs to their potential target genes, whose expression may be dysregulated by these non-coding SNPs. Although there were no significant associations, it was found that using SNPs on non-coding genes (Table 4), NF-κB transcription factor activity pathway and protein complex assembly pathway were both identified associated with HP with small P-values. NF-κB is a transcription factor that induces expression of more than 200 genes involved in diverse process such as cell survival, cell adhesion, inflammation, differentiation and growth^32^. Activation of NF-κB up-regulates expression of its responsive genes in cancer cells including lung cancer cells^33,34^.

The major limitation of our study was the small sample size (24 cases and 24 controls) and the lack of replication using an independent sample. Future work is planned to both increase the sample size and replicate results in an independent sample. Given that HP is a rare disease we plan to use tissues from these patients to verify the pathways identified. As examples, verification of cell cycle activation could be performed thru immunolabeling of human lung specimens with antibodies to KI67 or PCNA^35^. To determine if the suggested activation of proteasomal function (Adjusted P=0.0009) is occurring in diseased human lung, immunolabeling of human lung biopsy specimens could be performed with antibodies against the UPR signaling molecules ATF4, PERK and phospho-IRE1^36^; these experiments are in progress.

